# Detailed Survey of an in-vitro Intestinal Epithelium Model by Single-Cell Transcriptomics

**DOI:** 10.1101/2023.05.23.541940

**Authors:** Ran Ran, Javier Munoz, Smrutiti Jena, Leopold N. Green, Douglas K. Brubaker

## Abstract

The gut plays a critical role in maintaining human health by facilitating the absorption of nutrients, regulating metabolism, and interacting with the immune system and gut microbiota. The co-culture of two human colorectal cancer cell lines, Caco-2 and HT29, on Transwell is commonly used as an *in vitro* gut mimic in studies of intestinal absorption pharmacokinetics, gut mechanics, and gut-microbe interplay given the similar morphology, expression of transporters and enzymes, and barrier function. However, to sufficiently evaluate the translatability of insights from such a system to human physiological contexts, a detailed survey of cell type heterogeneity in the system and a holistic comparison with human physiology are needed to be conducted rather than by the presence of a few well-studied proteins. Single-cell RNA sequencing provides high-resolution expression profiles of cells in the co-culture, enabling the heterogeneity to be characterized and the similarity to human epithelial cells to be evaluated. Transcriptional profiles of 16019 genes in 13784 cells were acquired and compared to human epithelial cells (GSE185224). We identified the intestinal stem cell-, transit amplifying-, enterocyte-, goblet cell-, and enteroendocrine-like cells together with differentiating HT29 cells in the system based on the expression of canonical markers in healthy adult human epithelial cells. The epithelium-like co-culture was fetal intestine-like, with less variety of gene expression compared to the human gut. Transporters for major types of substance (lipid, amino acid, ion, water, etc.) were found transcribed in the majority of the enterocytes-like cells in the system. However, some of the well-studied transporters such as FATP4 and GLUT2 were absent. Toll-like receptors were not highly expressed in the sample, yet the treatment of lipopolysaccharide still caused a mild change in trans-epithelial electrical resistance and gene expression, possibly by the interaction with CD14, the co-receptor for TLRs. Overall, the Caco-2/HT29 co-culture is a cost-effective epithelium model for drug permeability testing or mechanical simulation, but its phenotypic discrepancy with the real epithelium is not negligible. As a result, its response to biological factors might not provide transferrable knowledge to the study of human gut physiology, especially the innate immune aspect.

## Introduction

Inflammatory Bowel Diseases (IBD) are complex and debilitating diseases that affect millions of people worldwide^1^. These disorders are characterized by an exaggerated immune response that leads to chronic inflammation in the gastrointestinal tract, resulting in a range of symptoms including abdominal pain, diarrhea, and rectal bleeding^2^. The two most common forms of IBD are Crohn’s disease (CD), which can affect any part of the digestive tract and is characterized by transmural inflammation, and Ulcerative Colitis (UC), which is primarily confined to the colon and rectum and is characterized by continuous inflammation of the inner lining of the colon^2^. The prevalence of IBD and colorectal cancer in the United States has been increasing dramatically in recent years^3^. According to recent estimates, approximately 3 million adults in the US have been diagnosed with IBD, with an estimated annual healthcare cost of over $6 billion^3^. This scenario is exacerbated by the higher risk of IBD patients relapsing within 1 year (28.7%) and 2 years (38.4%) after therapeutic de-escalation^4^. Additionally, studies have shown that individuals with IBD have an increased risk of developing colorectal cancer, with the risk increasing with the duration and extent of the disease^5^. Clearly, IBD is a significant health problem that requires urgent attention. Understanding the biology of IBD and the interactions between the immune system, the epithelium, and the microbiome is critical for the development of effective therapies for this disease^2^.

Caco-2 cells, derived from human colorectal adenocarcinoma, have been extensively employed as a remarkable proxy for the intricate gut epithelium^6^. The salient features that render these cells an exemplary in vitro model include their ability to differentiate into enterocyte-like cells, recapitulating the intestinal absorptive characteristics^6^. Furthermore, Caco-2 cells exhibit brush border enzymes, tight junctions, and a plethora of nutrient transporters, providing an accurate reflection of the physiological environment^6^. Additionally, their robust and reproducible barrier function makes them indispensable for scrutinizing drug permeability and absorption studies^7^. Lastly, the amenability of Caco-2 cells to various experimental manipulations bolsters their utility in exploring the multifaceted interactions between the gut epithelium and dietary components, pathogens, or pharmaceutical agents^6^. On the other hand, HT29 cells, another human colon carcinoma cell line, have garnered significant attention as a valuable model for mimicking gut epithelial cells^8^. These versatile cells exhibit the ability to differentiate into diverse intestinal cell types, including goblet and enteroendocrine cells, thus providing an authentic representation of the intestinal cellular milieu^8^. Moreover, HT29 cells contribute to the understanding of mucus secretion dynamics, which play a critical role in maintaining gut barrier integrity and host defense^9^.

However, to sufficiently evaluate the translatability of insights from such a system to human physiological contexts, a detailed survey of cell type heterogeneity in the system and a holistic comparison with human physiology need to be conducted, not just by the presence of a few well-studied proteins^10^. Single-cell RNA sequencing (scRNA-seq) represents a cutting-edge approach to meticulously assess the resemblance between in vitro gut mimics, such as Caco-2 and HT29 cells, and the authentic human intestinal epithelium^11^. By offering a high-resolution transcriptomic landscape of individual cells, scRNA-seq facilitates the identification of distinct cell populations, thus enabling a comprehensive comparison of cellular composition and heterogeneity between these models and native tissue^11^. Furthermore, this powerful technique uncovers the dynamics of gene expression and operative signaling pathways in both systems, shedding light on their functional congruence^11^. Employing scRNA-seq in this context also aids in the discovery of potential limitations or disparities within in vitro models, fostering the refinement and optimization of these systems for enhanced physiological relevance^12^. As a result, the subsequent improvements in these models pave the way for more accurate and insightful investigations of gut epithelial interactions with external factors, ultimately advancing our understanding of human gastrointestinal physiology and disease^10^. In this study, we employed scRNA-seq to investigate the molecular signatures of the co-culture and assess its potential for producing results that can be generalized to the human gut.

## Result

### Determination of Human Cell Type-like Signatures Represented in the Germ-Free Co-culture Model

We cultured Caco-2 (ATCC, passage no. 8) and HT29 (ATCC, passage no.13) separately at 1x105 cells/ cm2 and seeded them together into the apical chamber of three Transwells (Corning Inc, Costar, USA) with a ratio of 9:1 on day 0. Two Transwells were used as controls, where cells were co-cultured and harvested on day 23. The other Transwell was treated with 1 ng/ml Escherichia coli O111:B4 lipopolysaccharides on day 22 and harvested on day 23. Then all samples were prepared for single-cell RNA sequencing as per the manufacturer’s instructions (10X genomics) (Fig. 1A). Cell morphology agreed with the previous description of the Caco-2/HT29 monolayers (Fig. S1A-B). The TEER values reported from control treatments were within the range of values reported by Hoffmann et al.^13^, indicating that the established Caco-2/HT29 co-culture system possesses properties comparable to those in other reported cases (Fig. S1C).

**Fig. 1:**
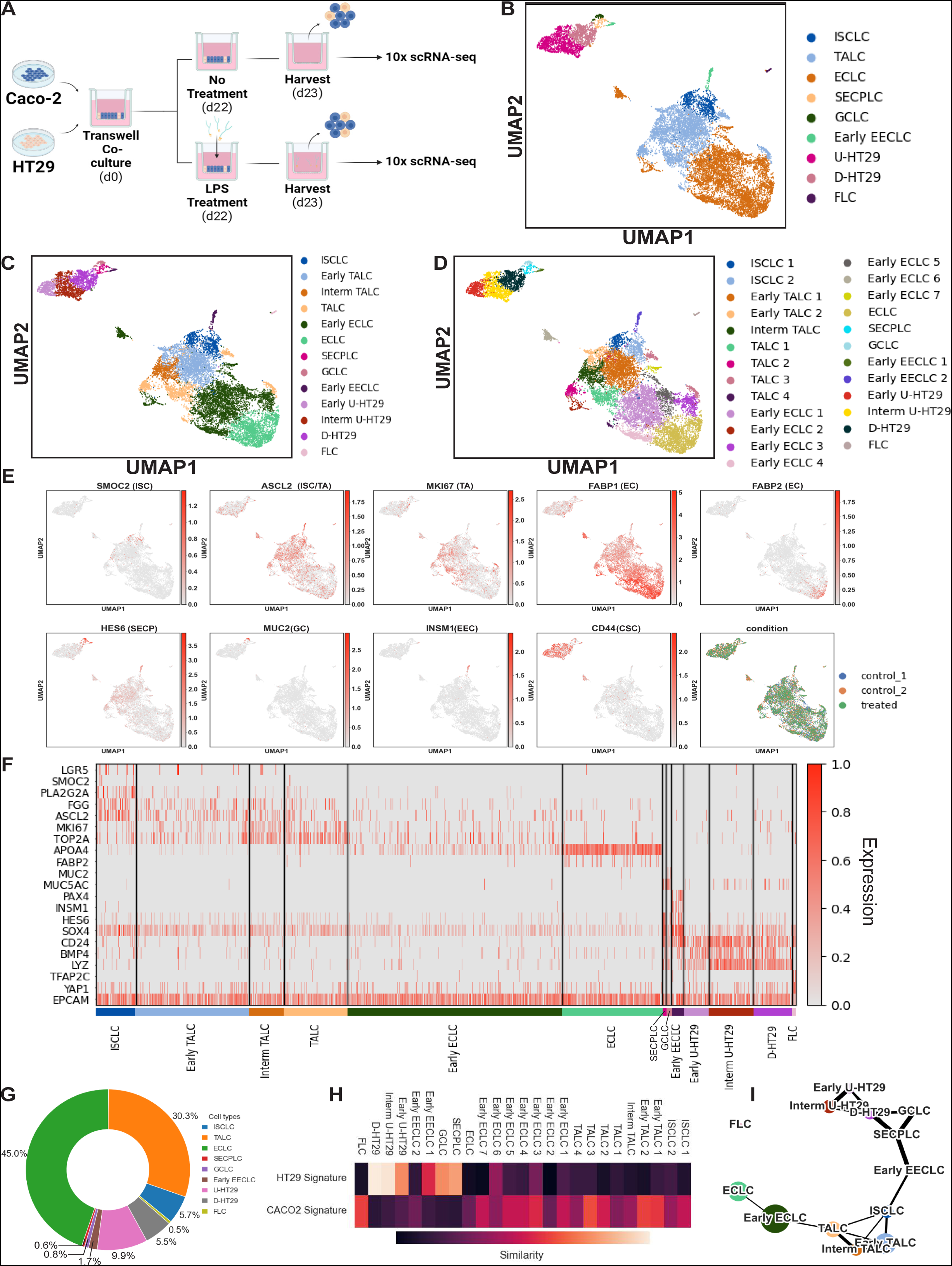
The Caco-2/HT29 co-culture model produces highly heterogeneous cell populations resembling intestinal epithelial lineage. A) Schematic of co-culture. B, C, D) uniform manifold approximation and projection (UMAP) of cell populations in the co-culture model with a resolution of (B) low (C) mid, and (D) high. ISCLC, intestinal stem-like cells; TALC, transit-amplifying-like cells; ECLC, enterocyte-like cells; SECPLC, secretory progenitor cell-like cells; GCLC, goblet cell-like cells; EECLC, enteroendocrine cell-like cells; U-HT29, undifferentiated HT29; D-HT29, differentiating HT29; FLC, fetal-like cells. E) UMAP visualization of co-cultured cells colored by key human intestinal epithelial lineage markers (cell types characterized by the markers are annotated next to the marker names) and by conditions (two controls, one LPS-treated group). F) heatmap of features in each cell population that have comparability with human intestinal epithelial lineages. G) composition of the co-cultured cells. H) conjectured Caco-2/HT29 origin of cell populations. Color bar) Jaccard similarity coefficient. I) Partition-based graph abstraction (PAGA) of the cell populations. Line thickness represents connectivity strength.

Cell identities were annotated based on their expression of previously reported signatures (ISC: LGR5, ASCL2, OLFM4, SMOC2, OLFM4^14^; TA: MKI67, TOP2A^15^; EC: APOA4, ANPEP, FABP2^16^; GC: MUC2^17^; EEC: PAX4, INSM1, CHGA, CHGB^18^) in healthy human intestinal epithelial cells (Fig. 1B, E). The general cell types were subdivided by their stemness/proliferating state (Fig. 1C) and were further segmented into more detailed cell groups (Fig. 1D) based on uniquely expressed genes (Fig. 1F). Cell subsets with no similar known human analogy were annotated based on their cancer-related gene expression and differentiation state. Overall, the Caco-2/HT29 co-culture system was highly heterogeneous. Quantitatively, about 6% of the co-cultured cells were intestinal stem cell-like cells (ISCLC), 30% were transit amplifying-like cells (TALC), 45% were enterocyte-like cells (ECLC), 1% were secretory progenitor-like cells, 2% were early enteroendocrine cell-like cells (EECLC), 1% were goblet cell-like cells (GCLC), and 15% were undifferentiated/differentiating HT29 cells (U-/D-HT29) (Fig. 1G). There was also a small cluster that exhibited fetal-like features, like the expression of TFAP2C (Table S1). All clusters had comparable sequencing depth (Fig. S2A) and mitochondrial gene percentages (Fig. S2B). The differentially expressed genes in each cluster were reported in Table S1. The conjectured origins of cell types, namely the monoculture Caco-2 or HT29, were determined by the overlap between cell type signatures and Caco-2/HT29 signatures (Table S2), identified from the bulk RNA-seq data of these two cell types provided by the Cancer Cell Line Encyclopedia (CCLE). Trajectory analysis of the system showed two distinct developmental branches, absorptive-like lineage ISCLC-TALC-ECLC and HT29/secretory-like lineages HT29-SECPLC-GC/Early ECLC (Fig. 1I). The arrangement of cluster nodes along the PAGA map followed the manually annotated time order.

We then asked how well the Caco-2/HT29 co-culture system can recapitulate the physiology of the human gut epithelium. We examined the expression profile of each major cell type in the model. Acknowledging that Caco-2 and HT29 were derived from colorectal adenocarcinoma, we also interrogated the cancer phenotypes of cells and asked if any of these properties disqualified the co-culture model for mimicking human gut epithelium.

### Characteristics of the stem and proliferative populations

Healthy human gut epithelium self-renews every 4-5 days. This quick turnover is initiated by the differentiation of ISCs located at the base of the crypt. Specifically, ISCs move upwards along the crypt-villi axis, leaving their stem niche and entering the region called transit-amplifying zone, where they rapidly proliferate and commit to either absorptive or secretory fate while transiting to the villus tip^19, 20^. In the co-culture, we identified 2 ISCLC subgroups, 2 early TALC subgroups, 1 intermediate TALC group, and 4 TALC subgroups (Fig. 2A). Noticeably, ISC and TA marker ASCL2, were universally expressed in all undifferentiated cells, which also agreed with the highly proliferative phenotype of cancer cells (Fig 2E-F, Fig. 2B).

**Fig. 2:**
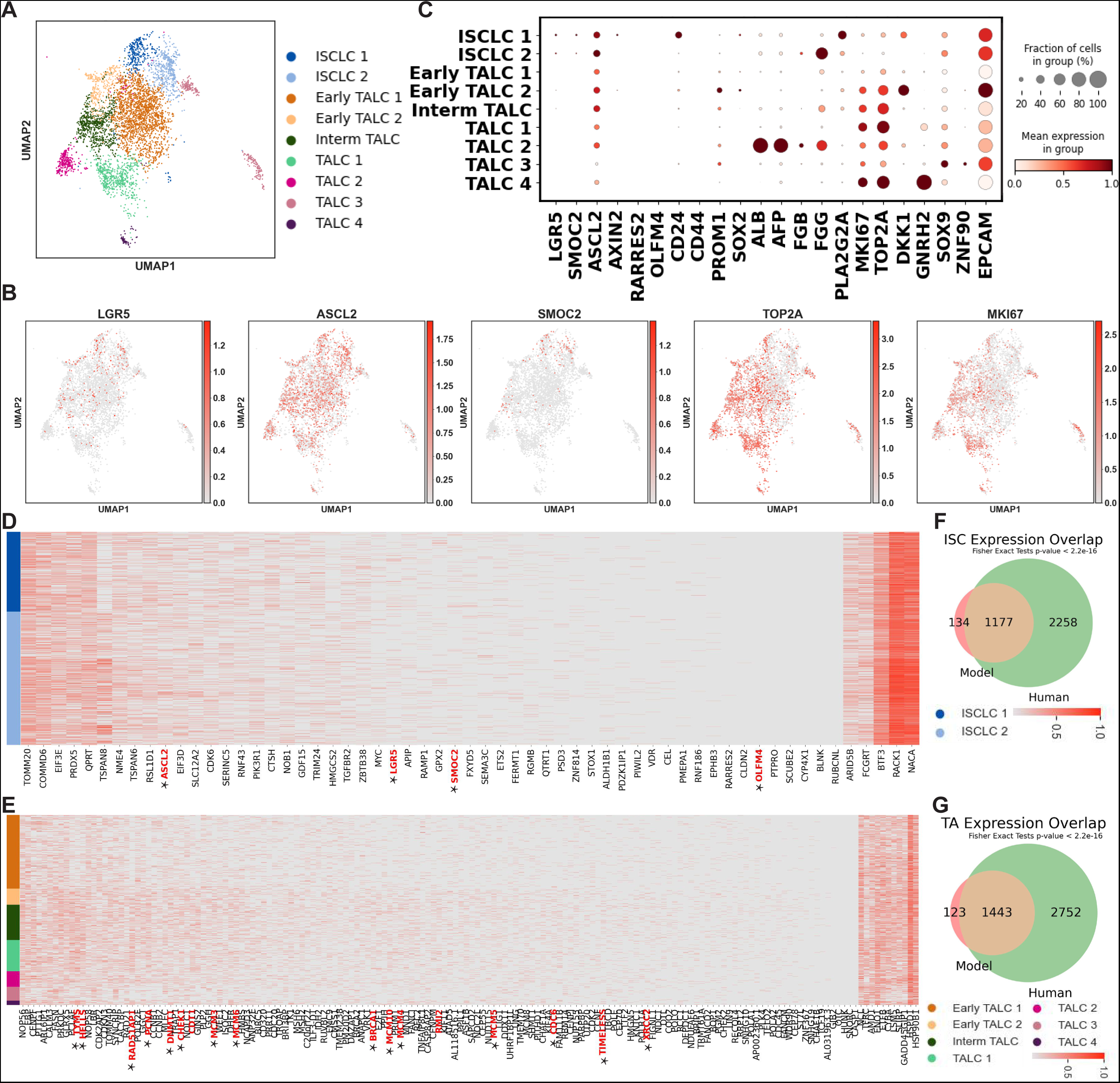
Characterization of ISCLC/TALC. A) UMAP of the ISCLC/TALC colored by detailed cell subtypes. B) Dot plot showing the featured gene expression in each ISCLC/TALC subtype. C) UMAP of the ISCLC/TALC colored by key human ISC/TA markers. D, F) Heatmap showing the expression of human (D) ISC markers in ISCLC and (F) TA markers in TALC. E), G) Venn diagram showing the overlap between commonly expressed genes (>30% of the cells) in E) ISCLC and human ISC G) TALC and human TA.

Both ISCLCs had a fraction of cells expressing classic intestinal stem markers LGR5 and SMOC2^14^ (Fig. 2B). A few cells from the ISCLCs were also expressing cancer stem cells (CSCs) marker PROM1 (CD133)^21^, fetal/cancer marker alpha-fetoprotein (AFP)^22, 23^ and its homologous albumin (ALB), and proliferative marker MKI67 and TOP2A at a relatively low level (Fig. 2C). ISCLC 1 uniquely expressed PLA2G2A, a phospholipase A2 family member that is highly expressed in fetal ISC^24^ and adult human Paneth cells (PC)^25^. PLA2G2A is known to be a Wnt inhibitor and PC differentiation modulator^26^, so its expression might indicate the Wnt activities and/or the identity of fetal small intestine ISCs or adult PCs. However, ISCLC 1 did not express PC markers LYZ^27^, DEFA4/5^27^, or PRSS2^28^ (Fig. S2C), so it was unlikely to be adult PC-like cells or the stem-like that have been reported to appear after inflammation in a previous study^29^. Given the ISCLC 1 also expressed ISC marker AXIN2^30^, CSC marker and embryonic development regulator CD24^31^, SOX2^32^, and BMP4^33^, it was most likely to be fetal ISCs with Wnt inhibition activities. By comparison, ISCLC 2 featured the strong expression of fetal ISC marker FGG while having little PLA2G2A. Since Wnt signaling maintains the self-renewal of ISCs^34^, the active transcription of the Wnt inhibitor PLA2G2A in ISCLC1 might indicate it was at the beginning of differentiation and cue the loss of stemness in this population while ISCLC2 was at a more undifferentiated state. Overall, the expression profiles of both ISCLC subgroups resembled those observed in fetal cells and cancer cells.

Two early TALCs had decreased ISC marker expressions and increased transcription of genes associated with proliferation such as MKI67 and TOP2B. Compared to early TALC-1, early TALC-2 still had a small fraction of cells that remained ISC/CSC markers positive. Meanwhile, this subgroup also highly expressed DKK1, a putative Wnt inhibitor^35^, and YAP’s downstream ANKRD1^36^. The potential Wnt inhibiting effect of DKK1 might signal the loss of stemness, thus agreed with the assigned TALC identity. In fact, the expression of DKK1 was negatively correlated with the stem regulator AGR2 and positively correlated with solute carrier family 2 member 3 SLC2A3 (GLUT3), indicating the decreased stemness and increased metabolic activities.

The activities of Wnt inhibitors in the stem/proliferative populations invite further investigation in the role of Wnt signaling pathway in the co-culture maintenance. We found the expression of genes that were engaged in the Wnt signaling, such as frizzled class receptor FZD 2/3/4/5/6/7^37^ and transcription factor TCF4^38^, as well as the transcription of several Wnt downstream genes like ASCL2, MYC^39^, ID1^40^, HES1^41^, SOX9^42^, etc., which means the co-cultured cells were universally eligible of receiving Wnt signals and proceeding the downstream transcription. Despite the broad expression of Wnt-associated genes, Wnt ligands themselves were not highly expressed. Only Wnt11 was transcribed in a faction of the co-cultured cells and Wnt10A in a few U-HT29 and D-HT29 cells at a low level (Fig. S3A). Such observation is reasonable, considering Paneth and mesenchymal cells are the main producers of Wnts in healthy human gut^43^, and the co-culture shares similarity with gut epithelium. Still, it raised the question how the Wnt pathway could be activated with such a limited Wnt secretion without the addition of exogenous Wnt (no Wnt protein in the culture medium or LPS treatment). It is possible that the secretion of Wnt11 activates non-canonical Wnt signaling^44^, the mutation on the downstream components such as APC/beta-catenin (which is well-characterized in both Caco-2 and HT29 cell lines^45, 46^), and/or the crosstalk between Wnt and other signaling pathways, such as TGF beta signaling^47^.

The ISC/CSC markers were further diminished in Intermediate TALC and TALC (Fig. 2B). TALC-1 featured high proliferative marker expression. The expression of AFP and ALB in TALC-2 was indicative of the resemblance of its features to the TA cells found in the fetal small intestine^22, 23^. This finding further supports the notion that the epithelium-like tissue derived from Caco-2/HT29 exhibits characteristics akin to the fetal intestine. Nevertheless, such finding was anticipated given the reported embryonic genes re-expressed or upregulated in colorectal adenocarcinoma^48^. TALC-4 also exhibited high levels of MKI67 and TOP2A, but uniquely expressed gonadotropin-releasing hormone 2 (GNRH2), a hormone with unclear function. Unlike GNRH1, GNRH2 is commonly expressed in a variety of human peripheral tissues but is seldom found in brain regions associated with gonadotropin secretion^49^, suggesting it might not function as a hormone in the same way GNRH1 does. Furthermore, it was observed that GNRH2 had an anti-proliferative effect on prostate, ovarian, breast, and endometrial cancer cells^50^. TALC-3 uniquely expressed Zinc Finger Protein 90 (ZNF90), a gene whose function is unknown but predicted to enable DNA-binding transcription factor activity^51^.

We examined the expression of previously reported human ISC/TA signatures in the model^14^. Although LGR5 and SMOC2 were expressed, there was no active transcription of human adult ISC marker genes OLFM4, RARRES2, or CD44 (Fig. 2B, Fig. 2D). Interestingly, the lack of OLFM4 is a feature of the immature fetal intestine, strengthening the fetal-like features of the co-culture^52^. However, we did not find any BEX5+ uniform progenitors^24^ that are reported to occur during the first trimester (Fig. S2C), suggesting the features were not early fetal-like. Overall, some of the human ISC differentially expressed genes (DEGs) were not expressed in the ISCLC, and conversely, only a few of the ISCLC DEGs were expressed in human ISC (Fig. S4A). A total of 1177 genes were expressed in over 30% of the population in both ISCLC and human ISC, whereas 134 genes expressed in ISCLC were not commonly expressed in human ISC. Additionally, 2258 genes commonly expressed in human ISC were not commonly expressed in ISCLC (Fig. 2D). By comparison, TALCs in the co-culture model shared more similarity with the human TA cells. Proliferative genes such as mini chromosome maintenance genes (MCMs) and cell division cycle (CDCs), which were differentially expressed in human TAs, were also prevalent in TALCs (Fig. 2F). All DEGs and 1443 out of 1566 commonly expressed genes found in TALCs were also expressed in human TAs (Fig. 2G, S4B).

### Enterocyte-like cells and their absorption preference

Enterocytes express a wide array of transporters, which play crucial roles in the absorption and secretion of nutrients, ions, and drugs, thereby maintaining the integrity of the intestinal barrier^53^. These transporters can be broadly classified into families such as ATP-binding cassette transporters, solute carrier transporters, and ion channels, each with distinct functions and substrate specificities. The activity of these transporters is tightly regulated and can be affected by various factors, including genetic variations, diet, and medication use. Caco-2/HT29 system is a popular high-throughput model to study the drug intestinal absorption. This system performs well in testing drugs that are absorbed passively, and a variety of transporters have also been identified on the surface of the Caco-2 cells^54^. The comprehensive profiling of transporters on Caco-2/HT29 remains incomplete due to the limited characterization methods available for the co-culture system during its establishment. Here, we identified 7 early ECLC populations and 1 mature ECLC population that resembles small intestine enterocytes (Fig. 3A). Their ECLC identity was defined by the expression of major mature enterocyte transporters APOA4, ANPEP, and FABP2^53, 55, 56^. The degree of their differentiation was annotated based on the expression of proliferative markers in these cells (Fig. 3B). We queried the expression of 9 categories of functional proteins that are present and responsible for digestion and absorption in healthy human enterocytes (Fig. 3C). The signature panel revealed unique features for each cell type, thus demonstrating the heterogeneity within the ECLC population. Adjacent to the fetal-like TALC-2 on UMAP, early ECLC-2 was actively transcribing AFP and ALB. Similarly, ECLC-4 featured a high expression of GNRH2. Early ECLC-6 exhibited undifferentiated HT29 features (i.e., LYZ, CD24, CD44, BMP4) at a lower level compared to the identified U-HT29 group (Fig. S2C) and was also positive for several SI enterocytes signature genes (Fig. 3B-C).

**Fig. 3:**
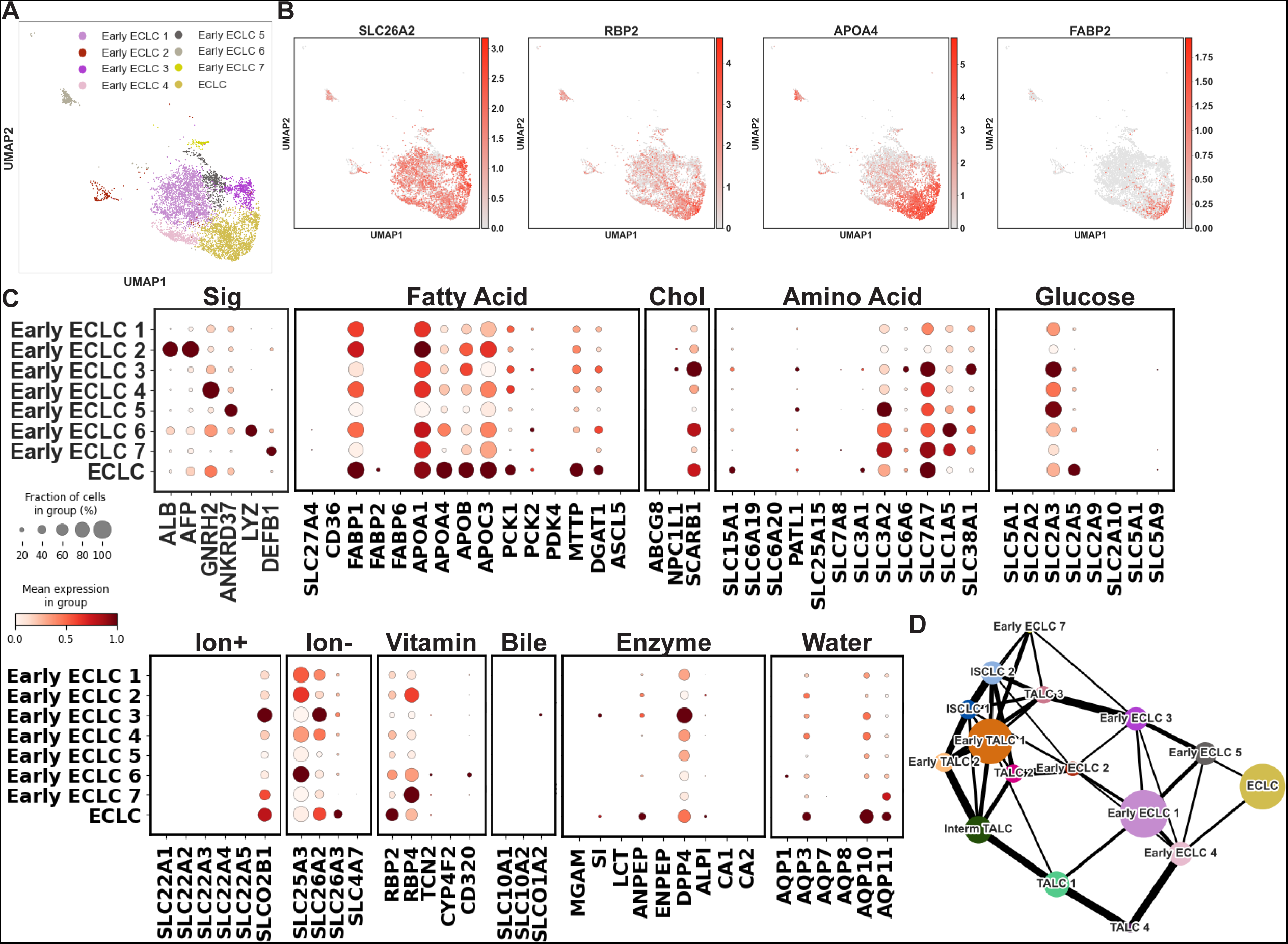
Characterization of ECLC. A) UMAP of the ECLC colored by detailed cell subtypes. B) UMAP of the ECLC colored by key human ECLC markers. C) Dot plot showing the expression of key transporters in each ELCL subtype. Sig, signatures; Chol, cholesterol. D) PAGA of the ECLC population. Line thickness represents connectivity strength.

Overall, all ECLCs expressed a reduced spectrum of transporters and enzymes compared to the human enterocytes. Observed in co-cultured cells were fatty acid transporters (APOA1/4, APOC3), digestive enzyme (ANPEP, DPP4), vitamin retinol/retinaldehyde metabolism RBP2/4, and water channel AQP3/10/11 (Fig. 3C). “They also actively transcribed several transporters including the drug transporter ABCC2, the bile acid transporter ABCC3, and the cation transporter SLCO2B1 (OATP-B). Notably, they also transcribed SLC2A3 (GLUT3), a glucose transporter whose activities are usually restricted to the brain but has also been detected in the placenta^57^ and CRC tissue^58^ (Fig. 3C). The GLUT3 activity increases the glycolytic capacity of cancer cells to fulfill their needs for rapid growth and metastasis^58, 59^. In the co-culture system, ECLC-3 and 5 expressed significantly higher GLUT3 than others.

However, the co-culture system lacked the expression of some well-studied essential transporters, such as SLC27A4^60^ (FATP4) or CD36^61^ for long-chain fatty acid absorption, ABCG5/G8 or NPC1L1 for cholesterol absorption^62^, SLC5A1 (SGLT1) or SLC2A2 (GLUT2) for glucose absorption^63^, ABCB1 for drug intake, lactase for enzyme digestion, SLCO1A2 (OATP3) or SLC10A2 (ASBT) for bile acid absorption^64^, AQP1, 7, or 8 for water transportation^65^, SLC22A1 through SLC22A5 (OCTs and OCTNs) or SCNN1B for cation transportation^66^. Taken together, we postulate that the metabolism of this group had been reprogrammed to involve certain cancer-related glycolysis burst events^67^, which skews the co-culture model toward cancer metabolism. Therefore, the performance of certain drugs, such as long-chain fatty acid drugs like omega-3 fatty acids, lipid-lowering drugs like Gemfibrozil/Bezafibrate, cholesterol-lowering drugs like statins/ezetimibe, glucose-lowering drugs like sulfonylureas, and other drugs targeting the missing receptors, might be inaccurately evaluated in drug screening or permeability/efficacy tests. Interestingly, previous study reported the existence of ASBT in Caco-2^68^, but we did not find its transcriptome in the co-culture.

While the model exhibited more small intestine (SI) characteristics (i.e., active SI-specific genes with most colon-specific genes missing), all absorptive populations expressed high levels of colon-specific ion transporter SLC26A2 and vitamin transporter RBP4 were expressed in all absorptive populations, indicating the model also possesses colon-like features given both Caco-2 and HT29 were CRC-derived. In unison, the heterogeneous ISCLC, TALC, and ECLC groups constitute a complex developmental roadmap (Fig. 3D).

### Undifferentiated HT29, Goblet cell-like cells, and immature enteroendocrine cell-like cells were present in the model

Undifferentiated HT29 cells, which are characterized by their rapid proliferation and ability to form a monolayer in culture without showing functional differentiation typical of mature enterocytes, such as the formation of microvilli and the expression of brush border enzymes. We found undifferentiated HT29 cells, secretory progenitor-like cells, goblet-like cells, and 2 groups of immature enteroendocrine-like cells present in the co-culture system (Fig. 4A). Previous reports have shown that the majority (∼95%) of HT29 monoculture is in an undifferentiated state, and they were defined such that they do not express definitive sets of markers of known human epithelial cell types^69, 70^. Admittedly, the profile of undifferentiated HT29, if any exists in the co-culture, is likely to differ substantially from that of the monoculture due to communication with Caco-2 cell and Transwell geometry. Therefore, we assigned the identities to U-HT29 and D-HT29 because of their lack of epithelial markers and share the most similarity with the profile of monoculture HT29 (Fig. 1H, Fig. 4B).

**Fig. 4:**
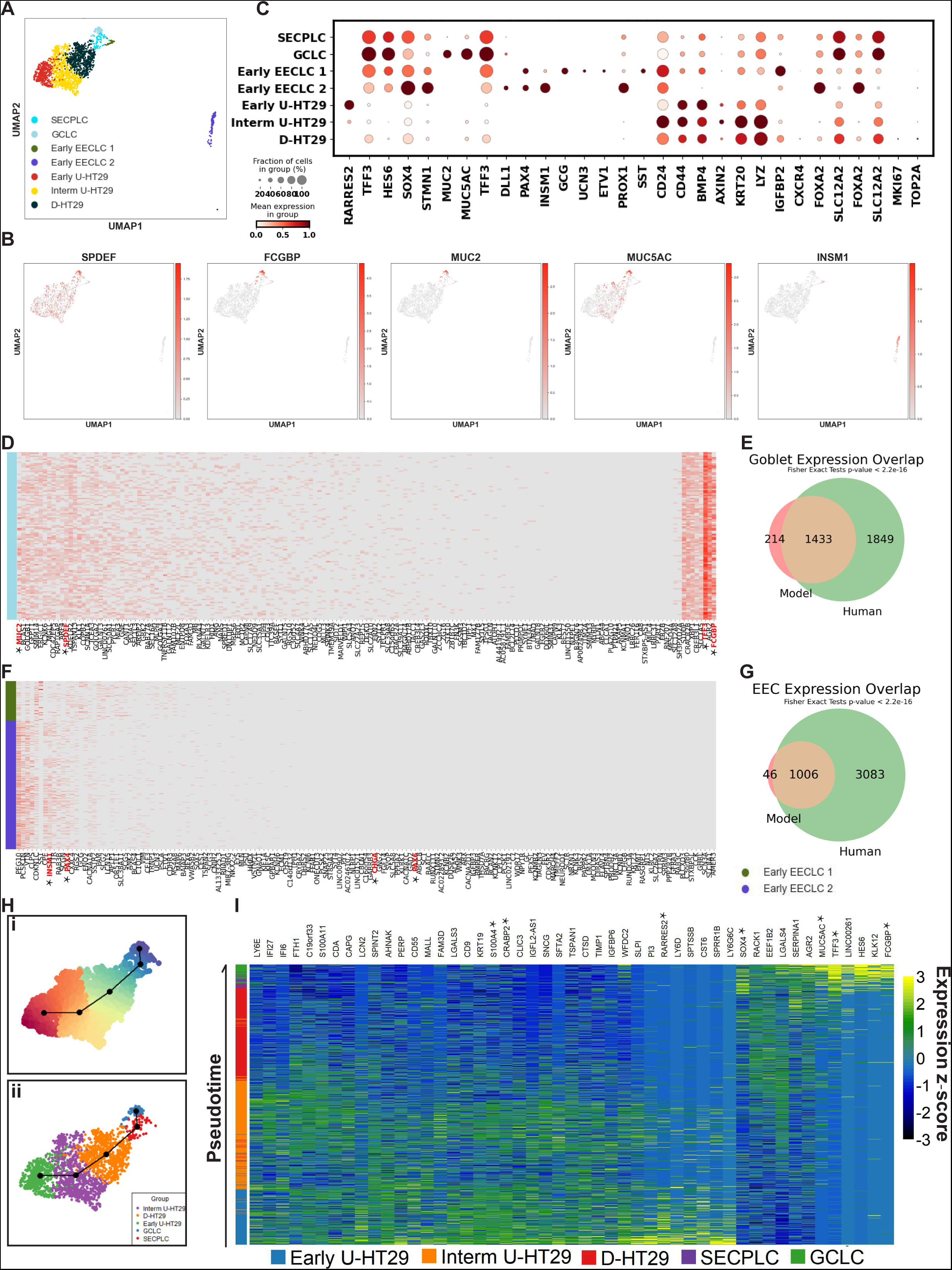
Characterization of secretory cell-like cells. A) UMAP of the HT29/secretory-like cells colored by detailed cell subtypes. B) Dot plot showing the featured gene expression in each HT29/secretory-like cells subtype. C) UMAP of the HT29/secretory-like cells colored by key human secretory (goblets/enteroendocrine) markers. D, F) Heatmap showing the expression of human (D) goblet markers in GCLC and (F) EEC markers in EECLC. E),G) Venn diagram showing the overlap between commonly expressed genes (>30% of the cells) in (E) GCLC and human goblet cells (G) EECLC and human EEC. H. UMAP of the HT29/secretory-like cells colored by (i) pseudotime, red = early, blue = late (ii) cell subtypes. I) Heatmap of gene expression changed along with the pseudotime. Expressions are standardized for each gene.

Most HT29 and secretory-like cells express CD24, CD44, AXIN2, and BMP4. CD24 and CD44, like discussed previously, are cell surface markers that are often used to identify cancer stem cells, which have the ability to self-renew and differentiate into various types of cancer cells. BMP4 is a bone morphogenetic protein that is reported to restrict the stemness of LGR5+ ISCs^71^ and enhance the differentiation^72^. Such a BMP4 expression evokes similarity to Paneth or mesenchymal stromal cells because the currently accepted gut epithelial cells turnover model proposes that a BMP4 gradient increasing from the crypt to the villus tip guides the differentiation of ISCs, and a high level of BMP4 is found in the lamina propria of the villus tip, co-localizing with LGR5^73^. However, the possibility of this population being stromal cells was undermined by the high expression of epithelial cell adhesion molecule (EPCAM). Interestingly, this population also highly expressed lysozyme (LYZ) and end-of-differentiation marker keratin 20 (KRT20)^74^. Lysozyme (LYZ), a well-studied antimicrobial agent, is transcribed in BEST4+ enterocytes, PCs, tuft cells, and follicle-associated epithelial cells, which seemingly supports the hypothesis that those cells being Paneth-like cells. Earlier studies also claimed HT29’s capability of differentiating into PCs based on its expression of LYZ^75^. However, the lack of other Paneth markers DEFA4/5 or ITLN2 (Fig. S2C) undermined the possibility that this population being full-fledged PC-like cells. KRT20 has long been used to identify the differentiated cells in the primary epithelium^74^, yet its role in CRC cells is controversial. Although it was found to be highly associated with well-differentiated CRCs, it was also evident to decrease as CRCs differentiate^76^. Furthermore, the RNA-seq of the monoculture undifferentiated HT29 provided by CCLE reveals a high expression of KRT20, which suggests that KRT20 might not work as a differentiated state indicator like in the gut epithelium context but an intrinsic property of cancer. SECPLC featured a high expression of fetal secretory marker HES6^23^, secretory differentiation modulator SOX4^77^, and microtubule regulator STMN1^78^ (Fig. 4B-C). It is also noteworthy that in the colorectal cancer context, these markers are associated with increased cell proliferation and invasion. The GCLC strongly and uniquely expressed gel-forming mucin MUC2^79^, a putative marker of the intestinal goblet cells and the building block of the mucus layer (Fig. 4B-C). It also highly expressed mucin MUC5AC^80^, which is the main component of the gastric and respiratory tract mucus layer that protects the tissue from stimulations yet is absent in the intestine^81^. Besides, part of the DEGs in human goblet cells were also present in the GCLC, such as IgGFc-binding protein FCGBP, SLC12A2, trefoil factor TFF3, and SPDEF (Fig. 4D). Although GCLC expressed some of the most well studied functional goblet genes, most of its the DEGs were not active in human goblet cells (Fig. S4C), suggesting the fundamental difference between healthy and malicious cells. Some of them, such as EGLN3^82^, ADAM8^83^, CA9^83^, AQP3^84^, and SIRT2^85^, are related to the hypoxia response, suggesting a hypoxia-related cancer metabolism in the GCLC. Specifically, CA9 is a well-known hypoxia-inducible gene that is commonly overexpressed in cancer cells^86^. Additional experiments, such as measuring oxygen levels in the cells would be needed to confirm if such a response is caused by the presence of hypoxia or the oncogenic signaling. Also, the intrinsic hypoxia genes expression means the Caco-2/HT29 mix might respond differently to oxygen levels or nutrient availability, so when using them as a gut epithelium mimic and co-culturing them with bacteria, their response might not be transferable to the real gut. Thus, it is essential to carefully consider the experimental conditions and controls to ensure that any observed effects are due to the bacterial co-culture and not to the Caco-2/HT29 cells’ intrinsic properties. When comparing with human GCs, we found that 1433 out of 1647 commonly expressed genes in GCLC were also prevalent in human GCs (Fig. 2G). Overall, the HT29-derived goblet cells were relative comparable to its healthy human epithelial cells’ analog.

Both early EECLC populations expressed immature EEC markers PAX4^87^ and INSM1^88^ (Fig. 4C). EECLC-1 uniquely expressed GCG, UCN3, ETV1 (EEC subtype L-cells), SST (EEC subtype D-cells)^23^, whereas EECLC-2 expressed PROX1 (EEC subtype L-cells) and a high-level of STMN1 (Fig. 4B). Most of the mature human EEC DEGs such as CHGA and CHGB were missing in both EECLC groups (Fig. 4F), yet some EECLC DEGs were expressed in human EECs (Fig. S4 D). While only 46 genes were uniquely expressed in either type of EECLC and not in human EECs, a significant number of genes (3083) present in human EECs were not universally found in EECLCs (Fig. 4G), which supported our inference that EECLCs in the co-culture system were immature (i.e., had a less variated expression profile than mature EECs). Taken together, EECLCs were found in co-culture, but they might not be a perfect in vitro model of mature EEC.

We further subdivided the undifferentiated HT29 population into early, intermediate, and late stages (also referred to as differentiating) and proposed a potential differentiation pathway from early U-HT29 to GCLC (Fig. 4H-I). This is based on the understanding that GCLC originates from HT29 and not from Caco-2 in a static culture^70^. Early U-HT29 uniquely and highly expressed oncogenic/metastasis genes CD9^89^, S100A4, and CRABP2^90^ (Fig. 4I). The level of classic stem markers LGR5 and SMOC2 was undetectable (Fig. S2C), yet it expressed ASCL2 and RARRES2, a gene transcribed mainly in colon ISCs and in colon goblet cells^56^ (Fig. 4B, Fig. 4I). The D-HT29 expressed metastasis markers insulin-like growth factor-binding protein IGFBP2^91^ and chemokine receptor CXCR4 (Fig. 4I, Fig. S2C). CXCR4 is often overexpressed in cancer cells and is known to play a role in cancer progression and metastasis^92^. Its signaling has been shown to promote cancer cell survival, migration, and invasion, as well as angiogenesis. In fact, the CXCR4 is known to be the marker of metastasis cancer stem cells^93^ (MCSCs) associated with the progression of CRC and a variety of cancers^94^. Other than IGFBP2 and CXCR4, the D-HT29 population also starts expressing secretory markers FOXA2^95^ and SLC12A2^96^ (Fig. 4B), indicating its secretory fate commitment. The intermediate U-HT29 cells didn’t exhibit any differentially expressed genes compared to early U-HT29 and D-HT29. However, their profile demonstrated a smooth transition between the two groups, which justifies our use of the term “intermediate” to describe their state.

### Co-culture model’s response to the E. coli LPS treatment

In the human epithelium, Lipopolysaccharide (LPS) triggers an immune response, like neutrophil activation, by binding to Toll-like receptors (TLRs) on epithelial cells^97^. TLR4 is primarily responsible for initiating the defense-related NF-κB signaling pathway in cells when it binds to the LPS of gram-negative bacteria^98^. Previous studies employ Caco-2/HT29 as models to study intestinal epithelial cells’ response to bacterial LPS by first confirming the presence of TLRs, or specifically TLR4. The undifferentiated HT29 is evident to express TLR4^99,100^, but that expression in Caco-2 is controversial. Some studies find the presence of TLR4 in Caco-2 using the western blot and report more intense TLR4 expression^100, 101^. Contradictorily, other studies report little TLR4 gene expression in Caco-2, with no observed LPS response^102–104^. This inconsistency is possibly due to the utilization of different anti-TLR4 and other technical differences during the cell culture. The bulk RNA-seq reads from CCLC support no TLR4 transcription in Caco-2.

In our co-culture system, neither TLR4 nor most other TLRs were detected in any sequenced and qualified cells under either condition (Fig. S5B). It is possible if 1) Caco-2, as reported, did not express TLR4, and 2) after the HT29 grew to confluence, its TLR4 expression dramatically decreased^102^.

Despite the lack of TLRs, we sought to understand how the system would change upon LPS treatment, given that another LPS-binding receptor, the glycosylphosphatidylinositol (GPI)- anchored protein (CD14), was present in the system, albeit at a very low level^105^ (Fig. S5C). Compared to the control, treated cells were less variate (Fig. 5A). The amount of early ECLC-6 (potentially HT29 derived cells), early EECLC-2 (potentially Caco-2 derived EECLC), ISCLC 2 (FGG+ ISCLC) was less than expected, whereas the Early ECLC-7 (DEFB1 high early ECLC) was more than expected (Fig. 5B). We then interrogated the differential expression after LPS treatment in ISCLC, TALC, and ECLC specifically, given that the purpose of the model is to gain insights of human intestinal epithelial cells (Fig. 5C). GCLC was excluded since statistical tests cannot perform well on a small-size sample. Overall, the transcription of tight junction genes TJP1, OCLN, and CLDN1/7 did not significantly differ between conditions (Fig. S5A). Inflammatory genes TNF-α, IL1β, and IFN-γ were not present/ did not change significantly upon the treatment (Fig. S5B). In ISCLC, genes like absorptive fate determination gene HES1 and fatty acid biosynthesis rate-limiting enzyme SCD were upregulated, while the fetal ISC marker FGG and TFF2/3 were downregulated (Fig. 5C(i)-D). In TALC, vitamin carrier retinol-binding protein RBP4 decreased, and CEACAM6, a receptor for adherent-invasive E. coli adhesion^106^, was upregulated (Fig. 5C(ii), E). SCD was also increased by the treatment in ECLC, while vitamin carrier retinol-binding protein RBP2 decreased (Fig. 5C(iii), F). Specifically, pathways related to differentiation, immune responses such as neutrophil activation, and metabolism were altered in the absorptive-like cells upon LPS treatment (Fig. 5G).

**Fig. 5:**
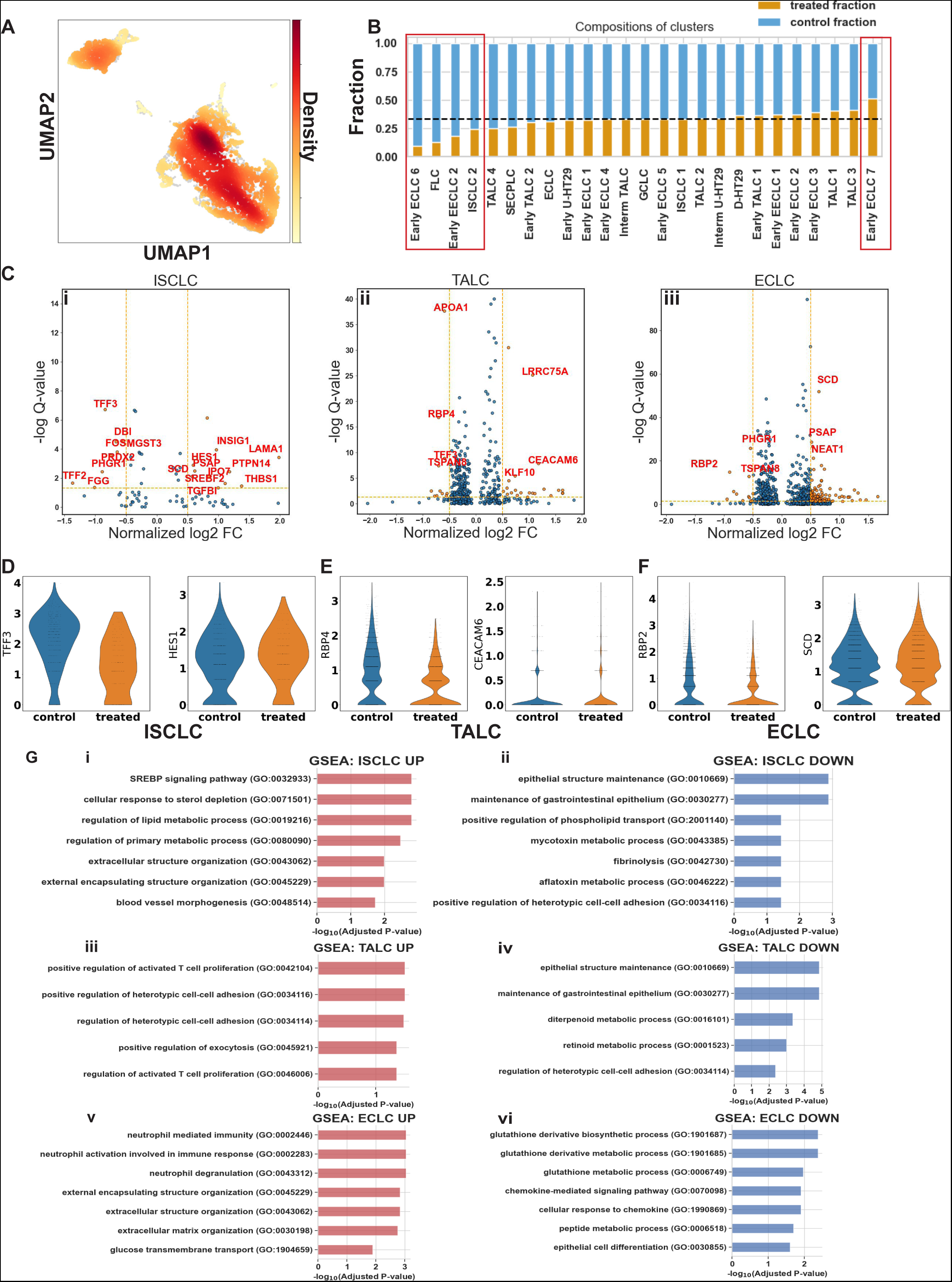
LPS treatment results in mild yet statistically significant expression change in ISCLC, TALC, and ECLC. A) UMAP of all the co-cultured cells, colored in the density of LPS-treated cells. B) Stacked bar plots showing the fraction of LPS-treated cells in each detailed cell population. Assuming the LPS treatment does not alter the composition of the system, the expected fraction taken up by the treated cells in each population should equal the total percentage of treated cells (indicated by the dashed horizontal line). C) volcano plots of differentially expressed genes in (i) ISCLC (ii) TALC (iii) ECLC across two conditions. FC, fold-change. Orange dashed horizontal line, -log Q > 2. Orange dashed vertical lines, log2 FC > 0.5 or < −0.5. Part of the DEs is annotated in red on the graph. D, E, F) Violin plots of one gene upregulated and one gene downregulated by LPS treatment in (D) ISCLC (E) TALC (F) ECLC. G) Gene set enrichment analysis (GSEA) result of pathways (i) upregulated (ii) downregulated in ISCLC, (iii) upregulated (iv) downregulated in TALC, (v) upregulated (vi) downregulated in ECLC.

## Discussion

The co-culture of Caco-2/HT29 has been widely used to simulate human intestinal epithelium in experiments. In our study, we carefully examined the variations in the Transwell co-culture, assessed how closely it resembled a healthy adult’s epithelium, and studied its response to E. coli LPS. We identified the ISC, TA, EC, GC, and EEC-like cells together with differentiating HT29 cells in the system based on the expression of canonical markers. Overall, the co-cultured cells showed less transcription variety and activities than the human epithelial cells, with a unique, intrinsic cancerous properties that confers the co-culture similarity towards the fetal gut given the common re-expression of embryonic genes in the colorectal adenocarcinoma. Earlier studies have compared Caco-2 and HT29 cells with the fetal colon, but due to previous limitations in transcriptomics and proteomics, only a limited number of genes were reported^107–109^. Another example of discrepancy is that enterocytes-like cells in the model did not fully express the major transporters in the small intestine or colon. However, to date, none of the “major transporters” has been proven indispensable for physiological intestinal fatty acid transport^110^. One possible explanation is that intestinal fatty acid absorption might be regulated by two or more genes so that one gene is inactivated, and the other could compensate for its function^110^. Therefore, although the transcriptome of some transporters is absent, substances might still be able to be transported. We also found no TLR4 and very marginal TLR2 expression in the co-culture. Although further experiments targeting the protein are necessary to confirm the absence of TLRs, here, we suggest that Caco-2/HT29 might not be an ideal model for studying microbe-epithelial cells interactions.

The Caco-2 and HT29 cell lines have proven extremely useful in replicating the gut epithelium, offering researchers a valuable laboratory model for exploring gastrointestinal physiology^111^. The Caco-2 cells, exhibiting enterocyte-like properties, primarily contribute to the understanding of absorptive processes, while the HT29 cells, characterized by mucus-secreting goblet cells, elucidate the protective mucosal barrier functions^46^. This powerful synergistic approach has not only accelerated advancements in the study of nutrient absorption and drug delivery but has also facilitated breakthroughs in understanding host-microbe interactions when immune cell presents^112^, the effects of cytokines and inflammatory mediators on barrier function and the expression of tight junction proteins^113^, the adherence and invasion of pathogenic bacteria^114^, and molecular mechanisms of colorectal cancer^115^, including the role of genetic mutations and the effects of potential therapeutic agents^116^.

Caco-2 and HT29 cell lines are widely recognized as useful lab models for studying gut lining. Still, traditional animal models are often used to give a comprehensive, lifelike system for studying gut biology and confirming results from lab models. It facilitates the study of complex interactions among host genetics, immune responses, the microbiome, and different organs ^117, 118^. However, the ethical considerations, costs, and potential species-specific differences limit its universal applicability. Alternative in vitro models, such as gut-on-a-chip, have emerged recently, presenting unique advantages and drawbacks. It offers a dynamic, microfluidic environment that incorporates fluid shear stress and mechanical strain, more accurately simulating the physiological conditions of the gut^119, 120^. Additionally, this platform allows for co-cultures of multiple cell types, including immune cells and microbes, thereby fostering a more comprehensive understanding of the intestinal ecosystem^121^. Compared to the primary-derived experimental methods like this, Caco-2 and HT29 cells exhibit a limited gene expression profile, fetal-like characteristics, and inherent cancer properties that might deviate from the in vivo gut epithelium. Nevertheless, their ease of use and cost-effectiveness still make them attractive for preliminary research.

Initially, we had 2 control groups and 2 LPS-treated groups. One of the LPS-treated groups was discarded due to its less-than-desirable sequencing quality. While this scope limits the scope of our analysis, we believe that our remaining treatment group, in combination with our controls, still provided valuable insights into our research question. We attempted to determine the origin of Caco-2/HT29 cells by analyzing the expression signatures we discovered from a comprehensive RNA sequence of Caco-2/HT29. However, we have to admit that during 22 days of co-culture, if the influences that two cell types exerted on each other were significant, then it is possible that cells developed convergently and acquired the pattern of the other type of cells so that their similarity with the mono-cultured cells is not indicative of their origins. Even so, as this study mainly focuses on describing the tissue similar to the epithelium formed after differentiation rather than the process of differentiation, we think this would be an interesting topic for future study.

## Method

### Germ-free coculture *in-vitro* Transwell gut model

The intestinal epithelial Transwell system was established by human colon carcinoma cell lines Caco-2 (passage no. 8) and HT29 (passage no. 13) (ATTC, United States) at 1x10^5^ cells/ cm^2^ on permeable polyester membrane Transwell inserts. The cells were cultured in DMEM medium with Glutamax (Gibco, United States), supplemented with 10% (v/v) FBS, and incubated in a humidified atmosphere (5% CO2, 37 °C) for 21 days to reach the full differentiation stage. Induction of barrier dysfunction in Caco-2 and HT-29 cell monolayers was induced in the apical chamber by 1 ng/ml *Escherichia coli O111:B4* lipopolysaccharides (LPS) stimulation on day 22 for 24 hours. On day 23, three Transwells were prepared for library construction following the manufacturer’s instruction. The platform used for sequencing was Illumina NovaSeq 6000 S4.

### Tight junction evaluation by transepithelial electrical resistance and the number of cells alive

To determine tight junction integrity, transepithelial electrical resistance (TEER) was measured by an epithelial volt-ohmmeter (EVOM2; WPI, Berlin, Germany) with STX2 Chopstick probes. Measurements were performed at each culture medium exchange according to the manufacturer’s instructions. The final TEER value was determined by subtracting the TEER measurement from a blank transwell insert and multiplying it by the cell culture surface.

### Single Cell RNA seq Data Pre-processing

The FASTQ files of the sequencing data were input into the Cell Ranger^122^ pipeline by 10x genomics to be filtered and mapped to the human genome GRCh38 to generate a count matrix that contains the number of detected transcriptomes of each gene for each cell. The downstream analysis was done with scanpy^123^ (v1.9.1) on Python and Seurat^124–127^ (v4.2.0) on R. Each dataset (control 1, control 2, and LPS-treated) was pre-processed separately. Low-quality cells were dropped based on their extremely low UMI counts, high mitochondrial gene counts, low ribosomal gene counts, and a low number of uniquely expressed genes (Table 1).

**Table 1.**
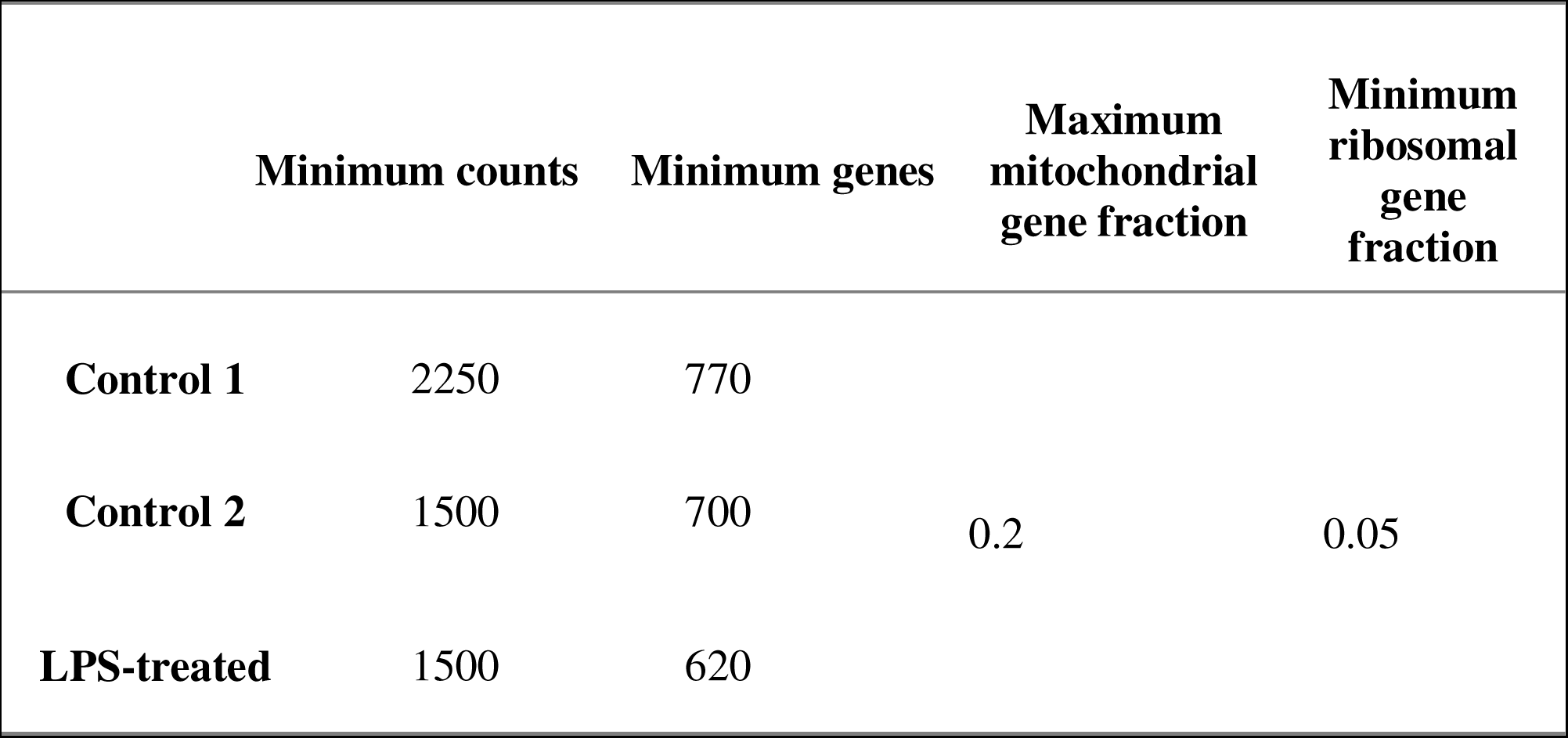
Pre-processing Parameters

Genes expressed in less than 5 cells in each dataset were excluded. Multiplets were removed by Scrublet^128^ (v0.2.3). After basic quality control, the 16019 genes’ transcriptional profile in 13784 cells was obtained. The cell cycle score was calculated based on the expression of cell cycle-related genes as previously described^129^. Each dataset was normalized by SCTransform^130^ (v0.3.5) on R. Cell’s mitochondrial fraction and the difference between the S phase score and the G2M phase score, as proposed as the representation of the cell cycle score, were regressed during the normalization. Then, the top 3000 genes that were most variable in as many datasets as possible were used as anchors to integrate the datasets in Seurat^131^. 3000 variable features were re-calculated for the combined dataset and used to perform the principal component analysis (PCA, 50 pcs). Leiden clustering^132^ (resolution = 0.65) was performed based on the computed neighborhood graph of observations (UMAP ^133^, 50 pcs, size of neighborhood equals 15 cells) to reveal the general subtypes of the co-cultured cells. Partition-based graph abstraction^134^ (PAGA) based on the Leiden clusters was used to initialize the uniform manifold approximation (UMAP, 50 pcs, min_dist = 0.1, spread = 1.1, n_components=2, alpha=1.0, gamma=1.0) to facilitate the convergence of manifold.

### Subclustering and cell identity labeling

The logarithmized SCTransform corrected counts were used to perform the Wilcoxon rank-sum test across every Leiden cluster to find the signature of a cluster, which were genes that have log2 fold change > 3 and adjusted p-value < 0.01. The resultant signatures, together with the epithelial markers reported previously, confirm the general identity of clusters. Each cluster was further subclustered to reveal more heterogeneity among cells. The resolution used in each subclustering is listed (Table 2). The Wilcoxon rank-sum test was re-run on the finer clustered data to unveil the marker of each subgroup, with the same threshold for log2 fold change and adjusted p-value.

**Table 2.**
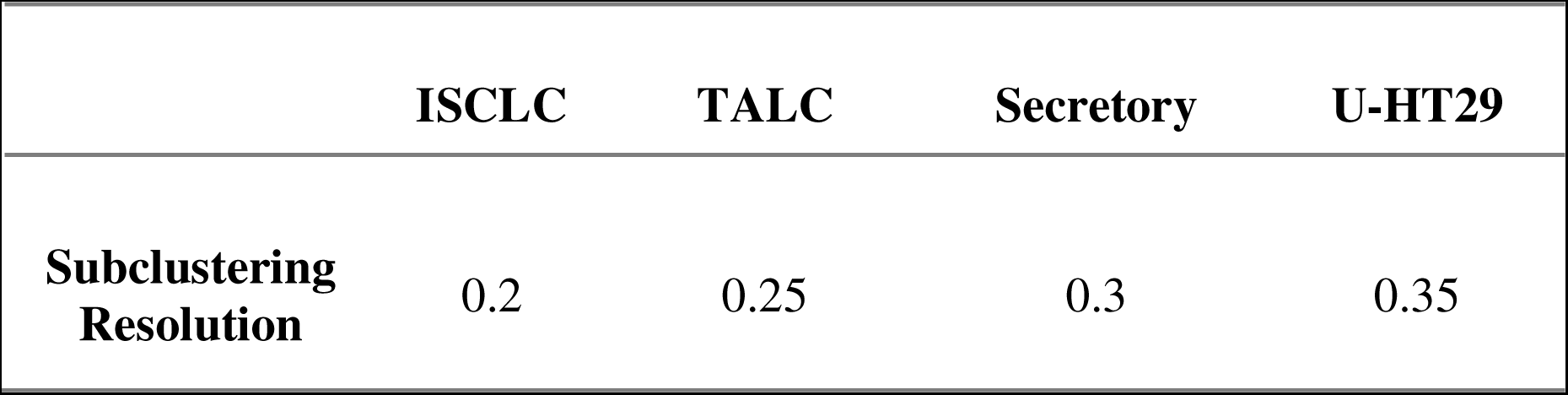
Leiden Subclustering Resolution

|  | ISCLC | TALC | Secretory | U-HT29 |
| --- | --- | --- | --- | --- |
| **Subclustering Resolution** | 0.2 | 0.25 | 0.3 | 0.35 |

### Caco-2/HT29 Similarity Determination

The raw bulk mRNA-seq counts of HT29 and Caco-2 cells were obtained from Cancer Cell Line Encyclopedia. Reads were rounded to integers. The positive and negative count differences between the two cell lines were fitted into two negative binomial distributions using statsmodel^135^ (v0.13.2), respectively. The overlap between the DEGs of each Leiden cluster and Caco-2/HT29 signatures were reported in Jaccard indices, normalized by the reference signatures size, and visualized in the heatmap.

### Differentially Expressed Genes Analysis

To determine genes that were differentially expressed in the same type of cells between control and LPS-treated, Monocle3^136–139^ (v1.2.9) was used to fit the SCTransform-corrected counts with a quasi-Poisson model to regress the gene expression over the treated/untreated condition. DE analysis was performed on treated/untreated ISCLC, TALC, and ECLC, given their adequate size for conducting statistical tests and their overall expression similarity to their human epithelial cells equivalent discussed earlier. The DE was defined as |log2 fold change| >= 0.5 with a Q value <= 0.05.

## Data Availability

The Caco-2/HT29 coculture’s scRNA-seq data can be accessed on NCBI GEO under GSE233628. Script used for analysis can be found on GitHub (https://github.com/WeldonSchool-BrubakerLab/Caco2-HT29-scRNAseq).

## Authors Contribution

Conceptualization: J.M., L.G., D.B.;

Methodology: R.R., J.M., S.J., D.B.;

Analysis: R.R.;

Writing–original draft: R.R., J.M.;

Writing–review & editing: R.R., J.M., S.J., L.G., D.B.;

Administration: D.B.

---

**Fig. S1:**
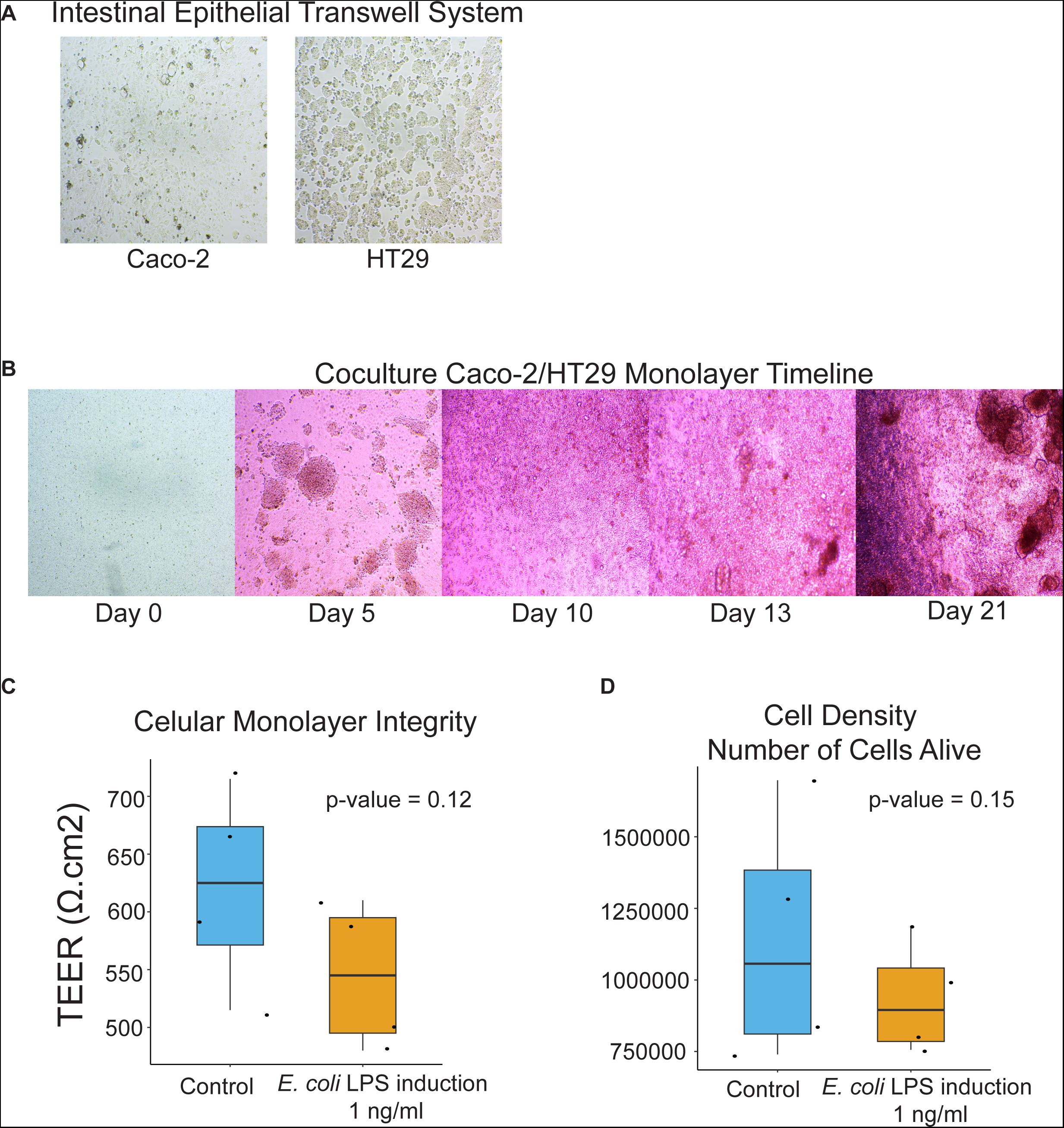
Physical properties evaluation of the co-culture. A) Brightfield microscopy shows the Caco-2 and HT29 monoculture morphology before co-culture. B) Brightfield microscopy shows the change in morphology and confluency during the co-culture. C) Trans-epithelial/endothelial electrical resistance (TEER) measurement with/without LPS treatment. Student t-test p = 0.12. D) Cell viability with/without LPS treatment. Student t-test p = 0.15.

**Fig. S2:**
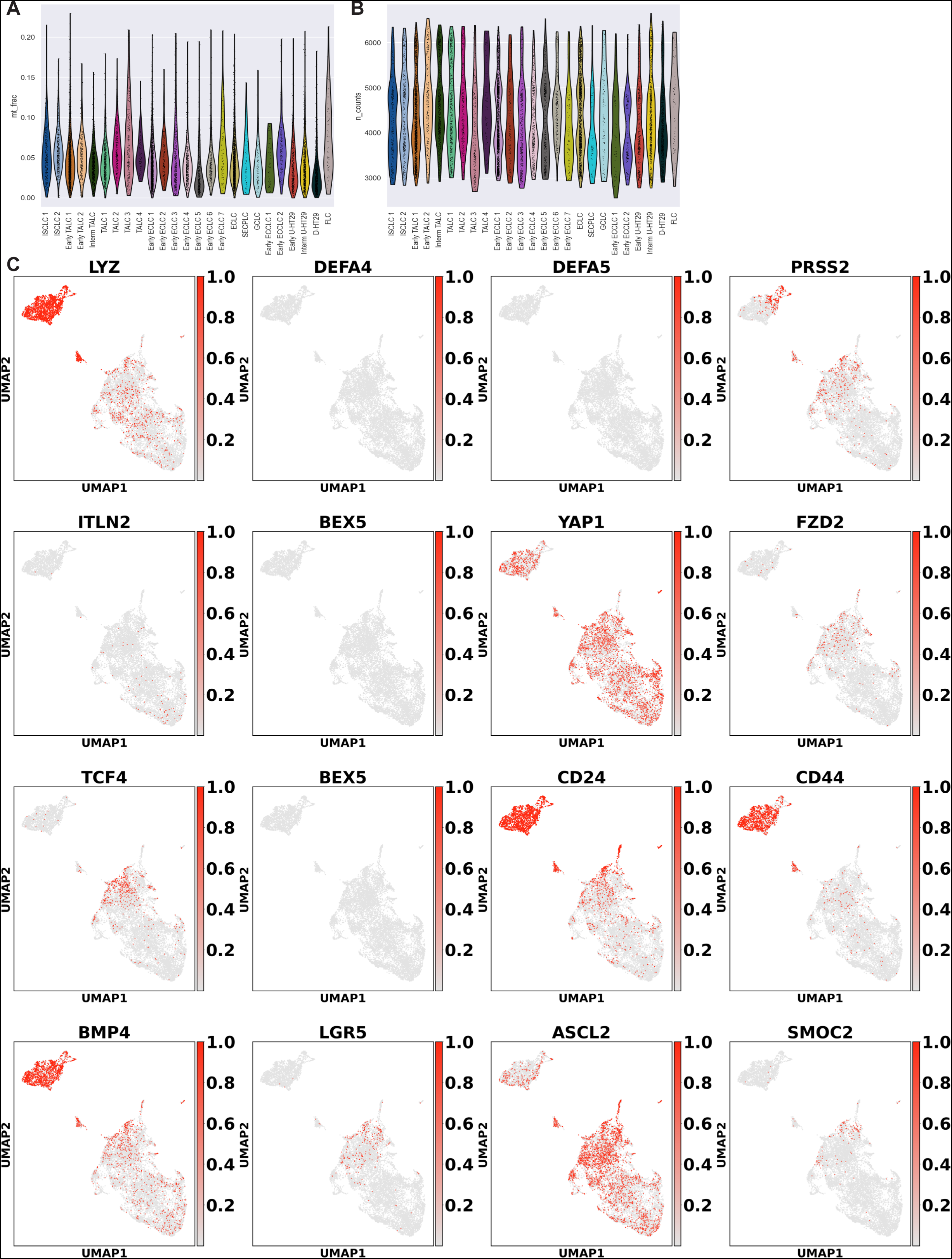
Quality control and supplementary expression maps. A) Violin plot of the mitochondrial fraction of each detailedly annotated cluster. B) Violin plot of the sequencing depth (i.e., counts of mRNA per cell) of each detailedly annotated cluster. C) Expression of various cell signatures on UMAP embedding.

**Fig. S3:**
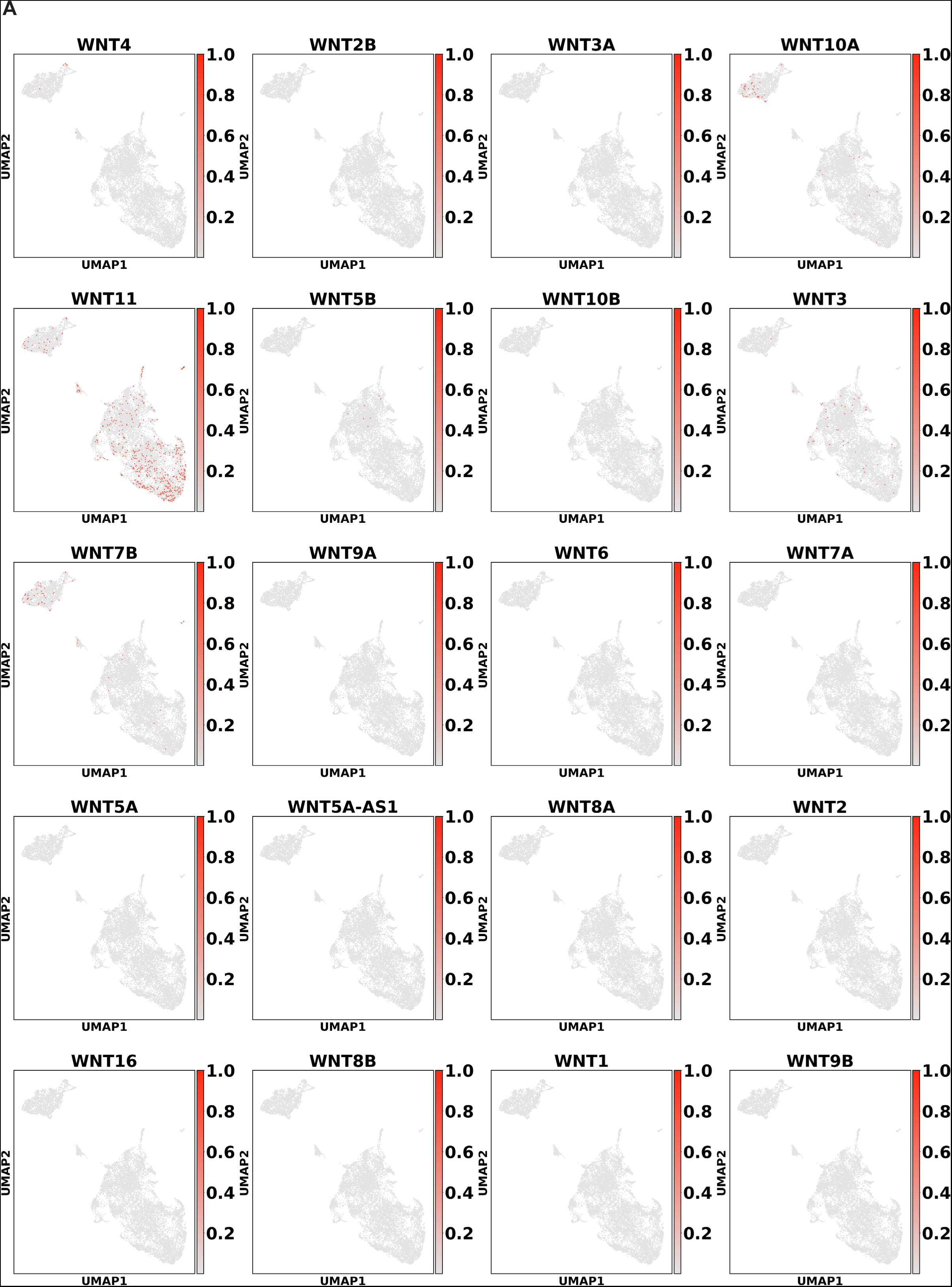
Expression maps of WNTs and CD14. A) Expression of WNTs and CD14 on UMAP embedding.

**Fig. S4:**
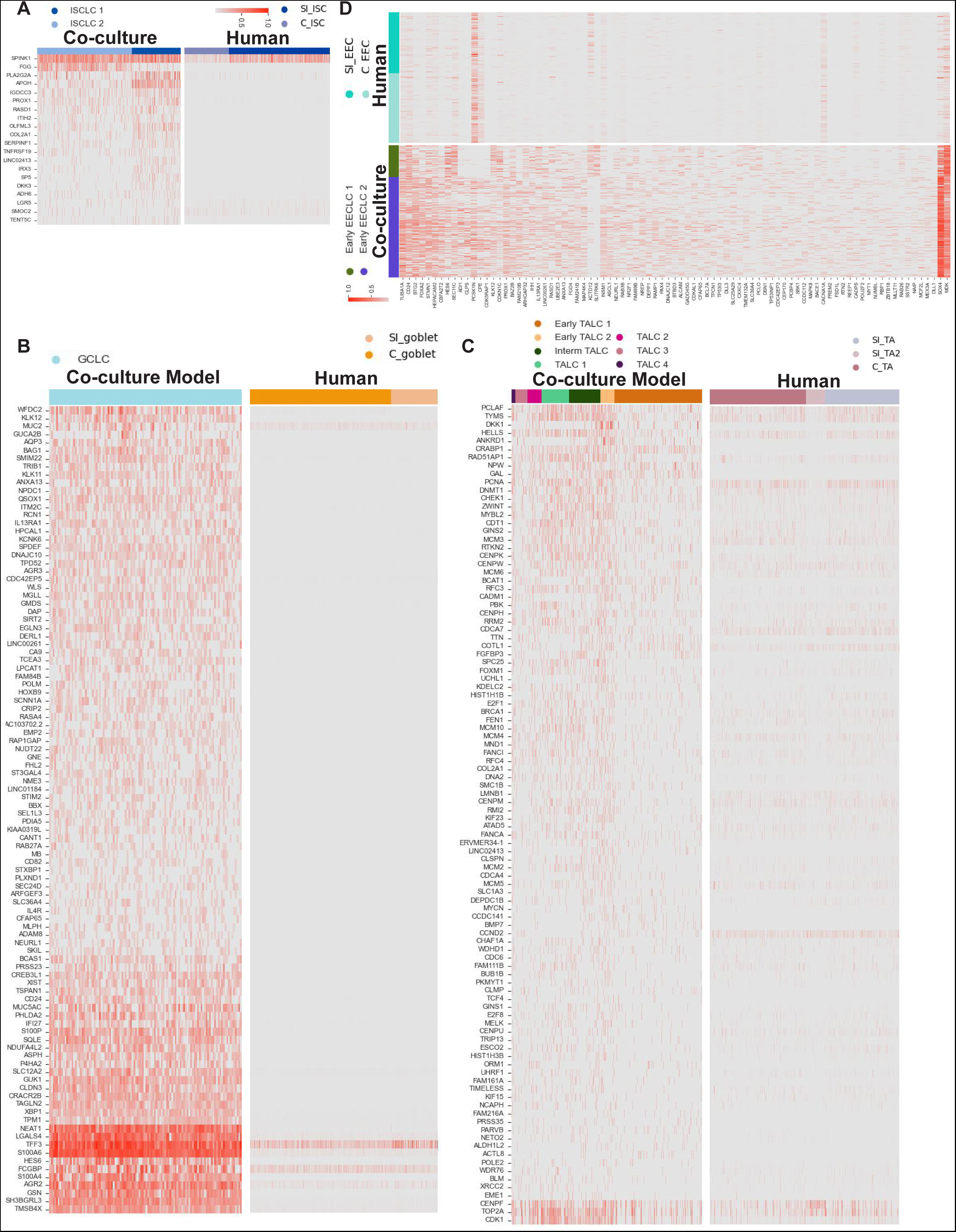
Heatmaps of model DEGs’ expression in corresponding human epithelial cells. (A)ISCLC DEGs in human ISCs. (B) TALC DEGs in human Tas. (C)GCLC DEGs in human GCs. (D)EECLC DEGs in human EECs.

**Fig. S5:**
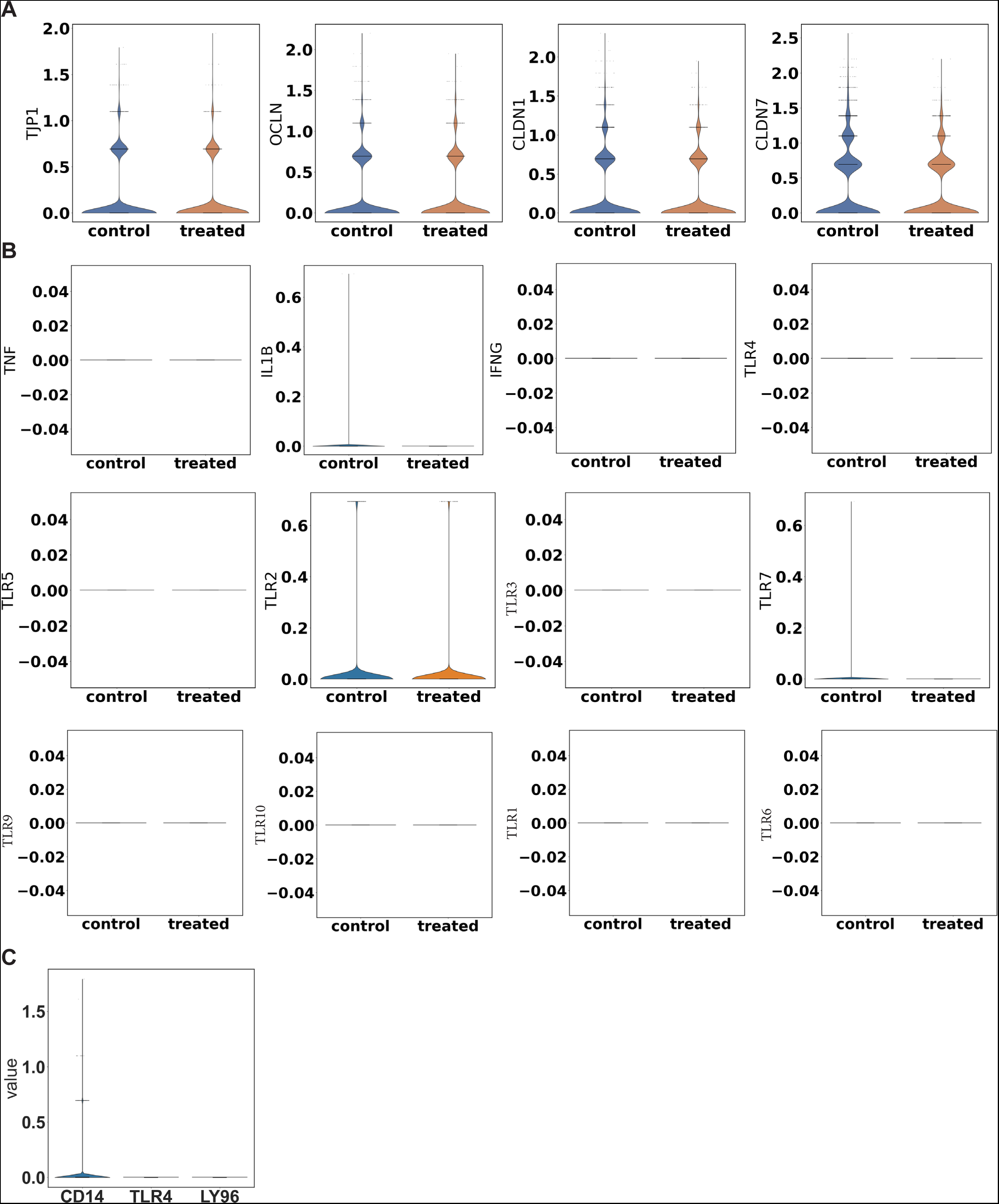
Comparison of gene expressions between control and treated conditions. A) Violin plots of 4 representative intestinal epithelial cells tight junction genes expression across conditions. B) Violin plots of 3 inflammatory genes and TLRs expressions across conditions. C) Violin plot of detected CD14, TLR4, and LY96 transcription in the co-culture.

